# Systematic analysis of goal-related movement sequences during maternal behavior in a female mouse model for Rett syndrome

**DOI:** 10.1101/2020.12.21.423671

**Authors:** Parker K. Stevenson, Devin M. Casenhiser, Keerthi Krishnan

## Abstract

Parenting is an ethologically relevant social behavior consisting of stereotypic components involving the care and nourishment of young. First-time rodent dams seek and gather wandering/scattered pups back to the nest (pup retrieval), an essential aspect of maternal care. Over the decades, qualitative observations of the behaving animal have been presented in quantitative discrete units. However, systematic analysis of the dynamic sequences of goal-related movements that comprise the entire behavioral sequence, which would be ultimately essential for understanding the underlying neurobiology, is usually not analyzed. Here, we present systematic analysis of pup retrieval behavior across three days in alloparental female mice (Surrogates or Sur) of two genotypes; *Mecp2*^*Heterozygotes*^ (Het), a female mouse model for a neuropsychiatric disorder called Rett syndrome and their wild type (WT) siblings. Additionally, we analyzed CBA/CaJ and C57BL/6J WT surrogates for within-strain comparisons. Frame-by-frame analysis over different phases was performed manually using DataVyu software.

We previously showed that Het are inefficient, by measuring latency and errors, at pup retrieval. Here, we show that the sequence of searching, pup-approach and good retrieval crystallizes over time for WT; this sequence does not crystallize in Het. We found that goal-related movements of Het in different phases were similar to WT, suggesting context-driven atypical dynamic patterns in Het. We also identified pup approach and pup grooming as atypical tactile interactions between pups and Het, which contribute to inefficient pup retrieval. Day-by-day analysis showed dynamic changes in goal-related movements in individual animals across genotypes and strains in response to the growing pups. Overall, our approach 1) embraces natural variation in individual mice on different days of pup retrieval behavior, 2) establishes a “gold-standard” manually curated dataset to next build behavioral repertoires using machine learning approaches, and 3) identifies distinct atypical tactile sensory processing in a female mouse model for Rett syndrome.

## Introduction

Maternal behavior is comprised of discrete units of behaviors, whose genesis and maintenance are differentially expressed throughout the early lifetime of the young. These units of behaviors may be classified as passive (not involving movement) or active (involving goal-related movements) (Stern, 1996). Active behaviors are comprised of pronurturant units, such as nursing, nest building/repair, licking of pups, and pup retrieval. Passive behaviors are components of nursing which may be initiated by the dam or pup. Traditionally, these behavioral studies were quantified using qualitative “spot checks”, observations without disturbing the nest and cage (of 15 minutes or lesser), and later studies with short intervals of disturbances to induce pup retrieval or other maternal behaviors. These studies paved the way for more recent optogenetics experiments to perturb neural circuitry and molecular/cellular analysis to determine genes and cell types involved in maternal behavior (Bendesky et al., 2017; Chen et al., 2019; Chong et al., 2020; Fang et al., 2018; Kohl et al., 2018; Krishnan et al., 2017; Lau et al., 2020a; Lau et al., 2020b; Li et al., 2019; Maynard et al., 2018; Moffitt et al., 2018; Niv et al., 2015; Tasaka et al., 2018; Wu et al., 2014). However, due to limited manpower and/or due to reductionist views to focus on particular behaviors, only certain aspects of the dynamic behavior were noted and discrete end-point analysis (such as time to retrieve pups, nesting and errors in retrieval) performed in most studies. However, it is clear from observing maternal behavior, that recognizing pup stimuli and performing purposeful movements is a dynamic process, and quite variable across individual mice, ages, strains, different litters and laboratories. With the recent explosion of pose estimation and unsupervised machine learning approaches (DeepLabCut, LEAP, MoSEQ and SimBA) becoming available to behavioral and systems neuroscientists, and calls to authentically integrate behavior into neuroscience questions, we believe a manual systematic curation of pup retrieval behaviors is essential to interpret and provide context for these new applications, especially for social behaviors (Datta et al., 2019; Gomez-Marin and Ghazanfar, 2019; Mathis et al.,, 2018; Nilsson et al., 2020; Pereira et al., 2019; Pereira et al., 2020; Wiltschko et al., 2020).

Pup retrieval involves processing of primary sensory cues to direct efficient searching and gathering of pups with goal-directed movements back to the nest (Beach and Jaynes, 1956; Lonstein et al., 2015; Stern, 1996). However, it is clear from observing maternal behavior, that this process is dynamic, involving activation of an innate neural circuit by sensory experience, and quite variable across mice strains, different litters and individuals (Krishnan et al., 2017; Curley and Champagne, 2016; Champagne et al., 2007; Stern and Mackinnon, 1978). The richness of this dynamic process is reduced to a few data points and has not been well explored by the current analysis methods.

Virgin female mice (naïve) with no previous maternal experience can execute efficient pup retrieval after exposure to pups, and after co-housing with a pregnant mouse and her pups (Van Hemel, 1973; Alsina-Llanes et al., 2015; Koch and Ehret, 1989; Krishnan et al., 2017). We have previously shown that adult female mice with deficient MECP2 expression are poor at pup retrieval (Krishnan et al., 2017). *Mecp2*^*Heterozygous*^ female mice are a model for a neuropsychiatric disorder called Rett syndrome that predominantly affects girls and women. Rett syndrome is characterized by early typical development, followed by developmental regression, and issues with sensory, cognitive and motor deficits throughout life. The causative gene, MECP2, is found on the X-chromosome, and is thought to regulate gene transcription, in response to neural activity and experience (Amir et al., 1999; Lewis et al., 1992; Zhou et al., 2006). In these patients and the female mouse models, random X-chromosome inactivation leads to mosaic expression of the wild type MECP2 protein in some cells and the mutant protein in others. Due to this mosaic expression and heterogeneity in MECP2 mutations, Rett syndrome patients exhibit variable syndromic features that change through the lifetime. Thus, adult female *Mecp2*^*Heterozygous*^ mice are an appropriate model to study the variation in phenotypes over physiological states and time.

Previously, we established pup retrieval behavior in an alloparental behavior paradigm as a robust model to study the individual variability and to discern the underlying cellular and neural mechanisms in the mosaic brain (Krishnan et al., 2017; Lau et al., 2020a; Lau et al., 2020b). We utilized latency, a measure of time taken to retrieve scattered pups to the nest, and errors, where the adult interacted with the pup but did not successfully retrieve to the nest, as measures to quantify retrieval behavior. We also noted remarkable individual variability and change in retrieval behavior over days. In this current study, we revisit the pup retrieval behavioral paradigm with in-depth analysis to characterize the dynamic behavioral repertoire these female mice display.

## Materials and Methods

### Animals

All behavioral experiments were performed in adult female mice (10-12 weeks old). They were bred and maintained on a 12/12 h light/dark cycle (lights on at 7 A.M.) and received food ad libitum. Genotypes used were CBA/CaJ, C57BL/6J, *Mecp2*^*Heterozygous*^ (B6.129P2(C)-*Mecp2*^tm1.1Bird^/J) and *Mecp2*^*WT*^-siblings (Guy et al., 2001) (The Jackson Laboratory). *Mecp2*^*Heterozygous*^ mice are considered pre-symptomatic at this age (Krishnan et al., 2017). All procedures were conducted in accordance with the National Institutes of Health’s Guide for the Care and Use of Laboratory Animals and approved by the University of Tennessee-Knoxville Institutional Animal Care and Use Committee.

### Pup retrieval behavior

Pup retrieval behavior was performed as previously described (Krishnan et al., 2017) (Figure 1A). Briefly, we housed two C57BL/6J (C57) virgin female mice, one *Mecp2*^*WT*^ (WT, black) and one *Mecp2*^*Heterozygous*^ (Het, red), with a first-time pregnant CBA/CaJ (CBA, orange) female beginning 3-5 days before birth. Upon cohousing, the two naïve mice are now termed ‘surrogates’ or “Sur”. Pup retrieval behavior started on the day the pups were born (postnatal day 0; D0) as follows for each adult Sur mouse (Figure 1B).

**Figure 1.**
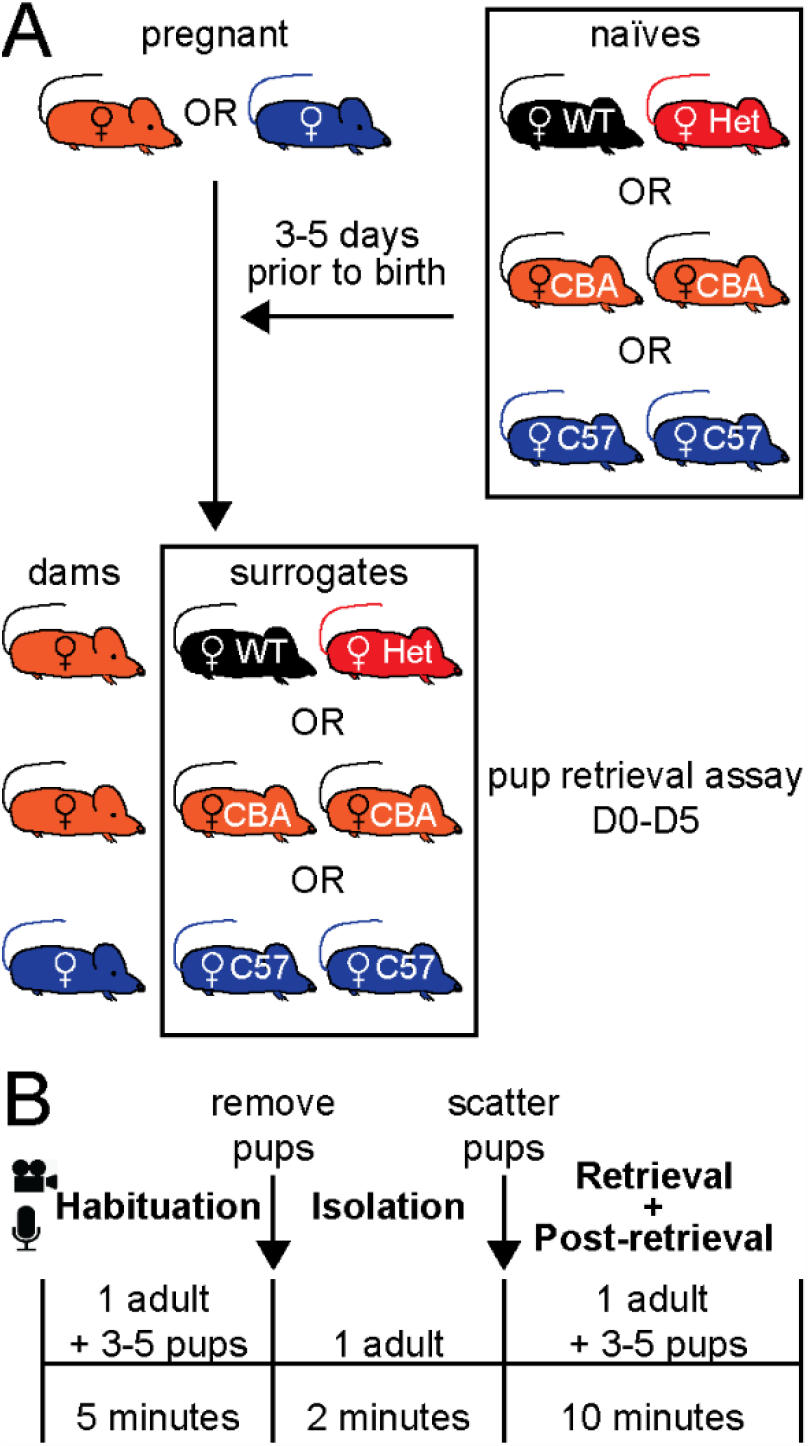
Schematic of behavioral analysis. **A)** Alloparental behavior setup. Naïve mice were added to home cage of the pregnant CBA (orange) or C57 (blue) mice 3-5 days prior to giving birth. The combinations were: 1) pregnant CBA mouse (orange) with naïve adult female C57 WT (black) and Het (red), 2) pregnant CBA mouse with 2 naïve WT CBA females (orange), and 3) pregnant C57 mouse (blue) with 2 naïve WT C57 (blue) mice. Pup retrieval behavior was performed on the day the mother gave birth (day 0, D0) and repeated daily up to day 5 (D5). Naïve mice are termed surrogates once the mother gives birth. **B)** Pup retrieval behavior setup. The two adults and any extra pups were removed from the home-cage and the remaining surrogate mouse was left undisturbed with 3-5 pups in the home cage for 5 minutes (Habituation). Then, pups were removed for 2 minutes (Isolation). Pups were scattered in home-cage and the adult allowed to retrieve and interact with pups for 10 minutes (Retrieval + Post-retrieval). All behavior were done in a dark box, and video 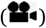- and audio 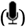-recorded.

1. Habituation phase - one adult mouse was habituated with 3-5 pups in the home cage for 5 minutes
2. Isolation phase - pups were removed from the cage for 2 minutes
3. Retrieval phase - pups were returned to home cage, one placed at each corner and at the center (the nest was left empty if there were fewer than 5 pups). Each adult female had maximum of 10 minutes to gather the pups to the original nest. Most WT retrieved all pups in less than 2 minutes.
4. Post-retrieval phase – time from when the last pup was retrieved to the end of the 10-minute recording session.

After testing, all adults and pups were placed in their home cage. The same procedure was performed again daily till D5. All behaviors were performed in the dark, during the light cycle (between 9AM and 6PM). Video (infrared; 30 frames per second) and audio were recorded. Additionally, as we used two different genetic strains in the above design, CBA dams and C57 surrogates, we performed similar experiments within strain (Figure 1A; CBA pregnant female with CBA surrogate WT (orange); C57 pregnant female with C57 surrogate WT (blue)), in order to account for strain-specific behaviors during pup retrieval (Brown et al., 1999; Champagne et al., 2007). In total, we analyzed six cohorts of WT and Het, 2 cohorts each of CBA and C57 surrogates (for within strain comparisons). The videos were blinded for further analysis.

Normalized latency was calculated using the following formula:

latency index = [(t_1_-t_0_) + (t_2_-t_0_) + … + (t_n_ – t_0_)]/n*L

where n = # of pups outside of nest

t_0_ = start of trial

t_n_ = time of nth pup gathered L = trial length

An error was scored when the Sur interacted with the pups (licking, sniffing, etc.) but did not successfully gather them to the nest or to another location in the cage.

### Behavior Coding

We used DataVyu software to curate/code the behaviors with frame-by-frame resolution (Datavyu Team, 2014). Datavyu produces a coding spreadsheet that allows labeling of behaviors and the duration through timestamps denoting the beginning and end of each behavior. The behavior codes for goal-related movements (GRM) are as follows:

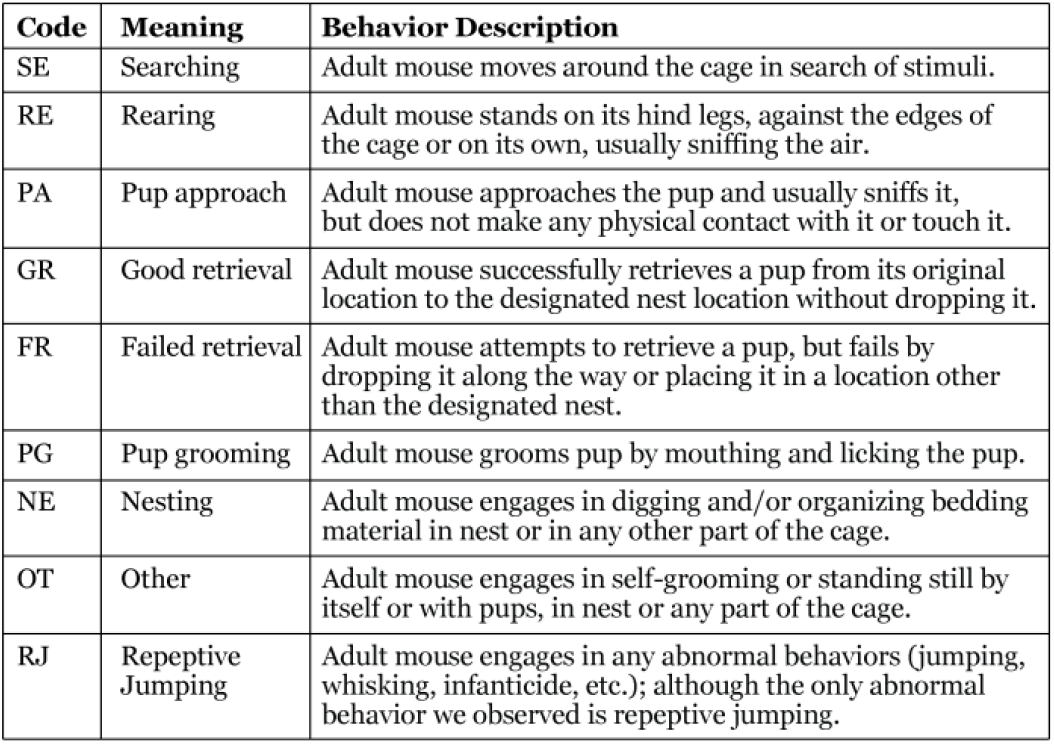

| Code | Meaning | Behavior Description |
| --- | --- | --- |
| SE | Searching | Adult mouse moves around the cage in search of stimuli. |
| RE | Rearing | Adult mouse stands on its hind legs, against the edges of the cage or on its own, usually sniffing the air. |
| PA | Pup approach | Adult mouse approaches the pup and usually sniffs it, but does not make any physical contact with it or touch it. |
| GR | Good retrieval | Adult mouse successfully retrieves a pup from its original location to the designated nest location without dropping it. |
| FR | Failed retrieval | Adult mouse attempts to retrieve a pup, but fails by dropping it along the way or placing it in a location other than the designated nest. |
| PG | Pup grooming | Adult mouse grooms pup by mouthing and licking the pup. |
| NE | Nesting | Adult mouse engages in digging and/or organizing bedding material in nest or in any other part of the cage. |
| OT | Other | Adult mouse engages in self-grooming or standing still by itself or with pups, in nest or any part of the cage. |
| RJ | Repetitive Jumping | Adult mouse engages in any abnormal behaviors (jumping, whisking, infanticide, etc.); although the only abnormal behavior we observed is repetitive jumping. |

### Analyses

Comparison of the percentage of time spent in each GRM was conducted using Unpaired Two-Samples Wilcoxon Tests in R version 4.0.2. Visualizations were produced using the *ggplot2* 3.3.2 and the *ggpirate* 0.1.2 packages (Wickham, 2016; Braginsky, 2020).

For Figures 5 and 6, GRMs were coded frame-by-frame during the 10-minute assay. Frames were compressed to 800ms bins for sequence plots display in the figure (Gabadinho et al., 2011). 1^st^ time bin marks the beginning of pup retrieval and ends at the 750^th^ bin (10-minutes). For Figures 7 and 8, transitional probabilities between GRMs were calculated as lag 1 discrete-time Markov chains with the aid of the *behavseq* package, and visualizations were produced using the *TraMineR* package versions 2.2 (Curley, 2015; https://github.com/jalapic/behavseq). All statistics and graphs were generated by using either R (2020) or GraphPad Prism (version 9).

**Figure 2.**
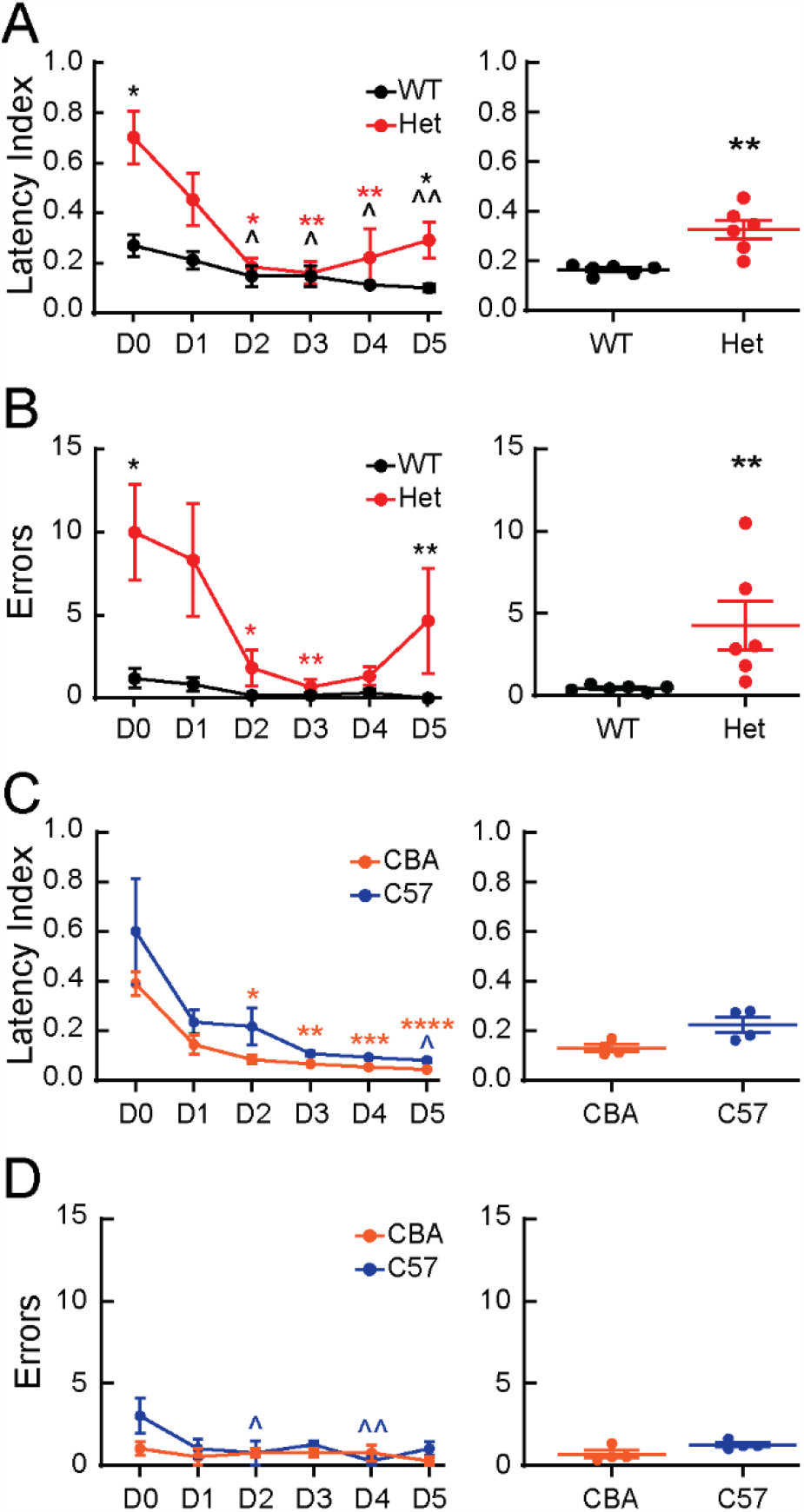
Pup retrieval performance of WT in two strains and *Mecp2*^*Het*^ (Het) across days: **A-B) Left:** WT mice retrieved significantly faster on D2-D5 compared to D0, as measured by latency index (A, *\*p* < *0*.*05, **p* < *0*.*01*), and made few errors in all 6 days (B). Compared to WT, Het retrieved significantly slower (A, *\*p* < *0*.*05*) and made more errors (B, *\*p* < *0*.*05, **p* < *0*.*01*) on D0 and D5. Compared to D0, Het showed significant behavioral improvement on D2-4 (A-B, *\*p* < *0*.*05, **p* < *0*.*01*) and was indistinguishable from WT. **Right:** Average of all 6 days revealed Het retrieved significantly slower (A) and made more errors (B) compared to WT (*\*\*p* < *0*.*01*). N = 5-6 animals per genotype per day. **C-D) Left:** Both WT CBA and C57 mice showed behavioral improvement by retrieving pups faster throughout the 6-day test period compared to D0 (C, CBA D0 vs D#: \**p* < *0*.*05*, \*\**p* < *0*.*01*, \*\*\**p* < *0*.*001*; C57 D0 vs D5: *\*p* < *0*.*05*). CBA and C57 also made fewer errors, with minimal but significant improvement on D2 and D4 by C57 compared to D0 (D, *\*p* < *0*.*05, **p* < *0*.*01*). **Right:** on average, CBA behavioral performance was similar to C57 by latency index (C) and errors (D). N = 4 animals per genotype per day. For A-D, mean ± s.e.m. are shown. *Statistics: Mann-Whitney test or Kruskal-Wallis followed by Dunn’s*.

**Figure 3.**
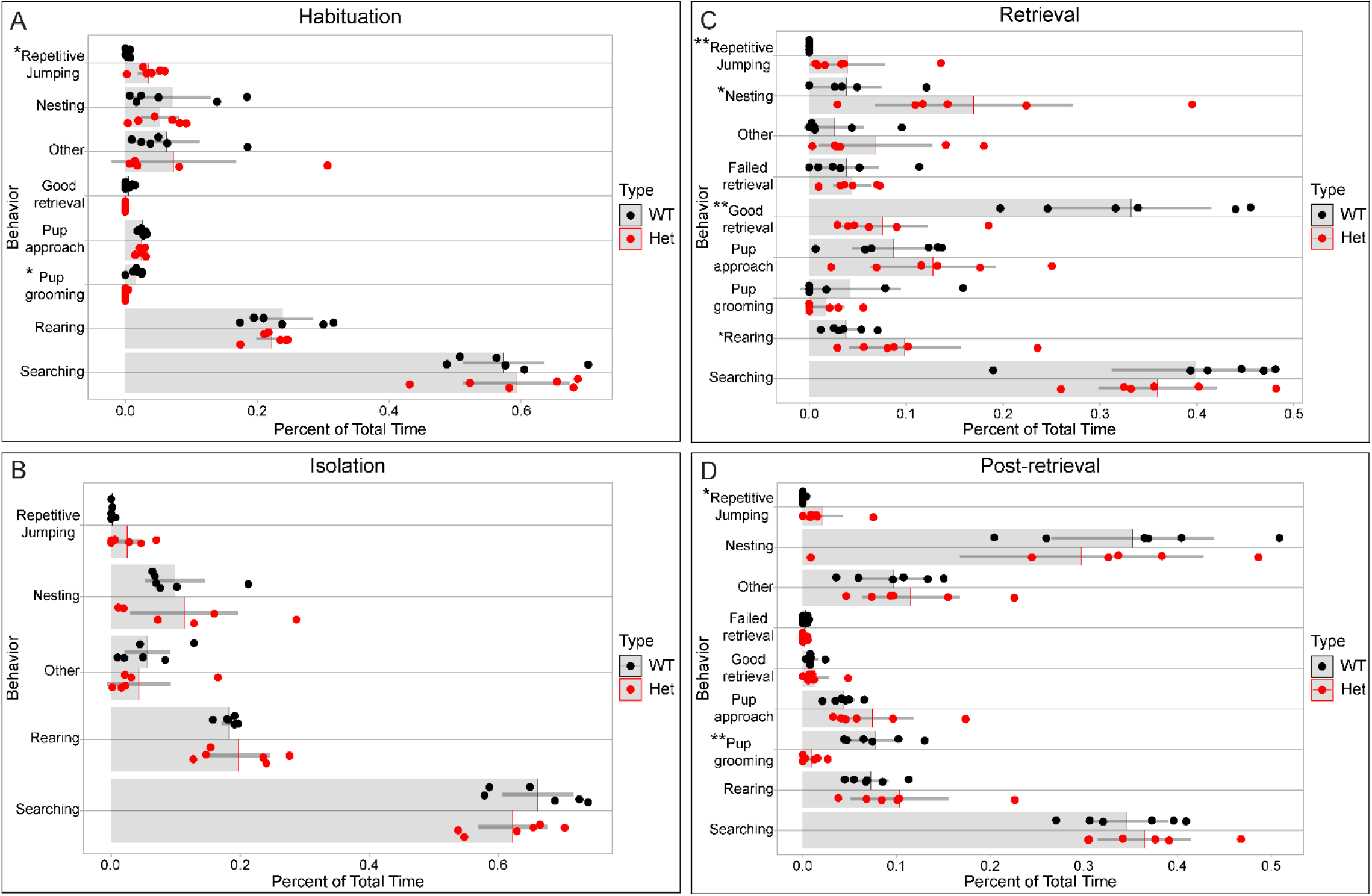
Het spend atypically more time in specific goal-related movements (GRMs) in the presence of pups. **A-D)** Compared to WT (black), Het (red) spent significantly more time with repetitive jumping during habituation (A), retrieval (C) and post-retrieval (D) but not in isolation (B). Het also spent significantly lesser time grooming pups in Habituation (A) and Post-retrieval (D). Additionally, during Retrieval (C), Hets spent significantly more time in nesting behavior and lesser time in good retrieval. Light grey bars indicate the mean proportion of time the mice spent in each GRM as a proportion of total time in all GRMs combined. Dark grey lines (whiskers) indicate 95% confidence intervals. Each dot represents the mean across 3 days for each mouse. *Statistics: Unpaired two-samples Wilcoxon test: *p* < *0*.*05, **p* < *0*.*01*.

**Figure 4.**
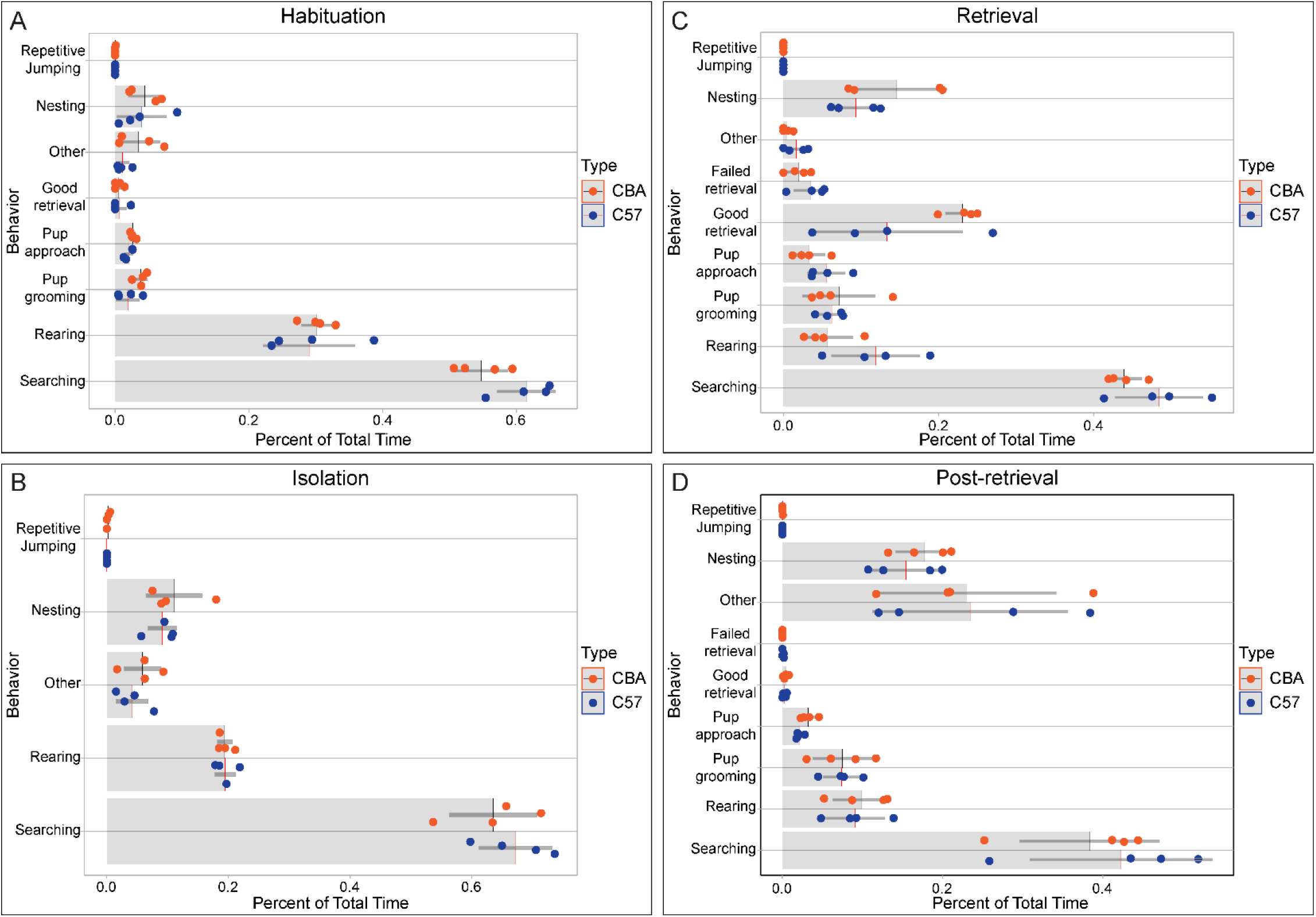
Across phases, no significant differences in time spent across GRMs in CBA and C57 WT. Habituation (A), Isolation (B), Retrieval and Post-retrieval phases (D). Light grey bars indicate the mean proportion of time mice spent in each GRM as a proportion of total time in all GRMs combined. Dark grey lines indicate 95% confidence intervals. Each dot represents the mean across 3 days for each mouse. *Unpaired two-samples Wilcoxon test: p* > *0*.*05*.

**Figure 5.**
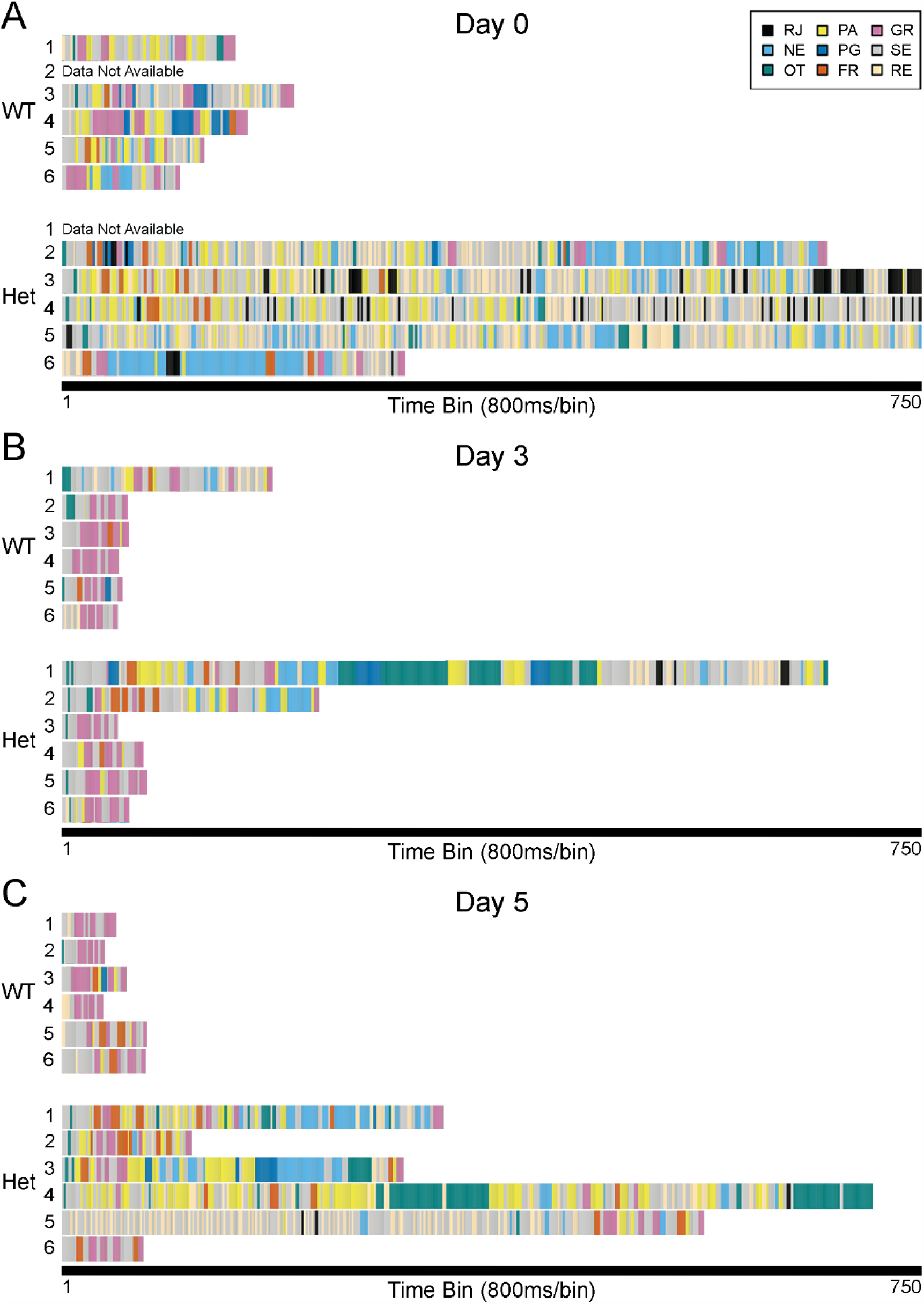
Natural variation in sequences of GRMs during retrieval across days in both WT and Het mice. **A-C)** While WT were comparatively efficient at pup retrieval on D0 (A, top) and maintained or improved their retrieval abilities on D3 (B) and D5 (C), Het were significantly less efficient and much more variable at retrieving pups (A-C, bottom). Interestingly, Het #2-6 show marked improvement on D3 but noticeably regressed on D5. GRMs were coded frame-by-frame during the 10-minute assay. Frames were compressed to 800ms bins for display in the figure. 1^st^ time bin marks the beginning of pup retrieval and ends at the 750^th^ bin (10-minutes). Each row depicts the sequence of behaviors for an individual mouse during retrieval phase. Mouse order is preserved across days to observe patterns and natural variation (example: mouse #1 is the same mouse on D0, D3 and D5). Abbreviations on codes (see Methods for details): RJ repetitive jumping (black), NE – nesting (light blue), OT – other (teal), PA – pup approach (yellow), PG – pup grooming (dark blue), FR – failed retrieval (orange), GR good retrieval (pink), SE – searching (grey), RE – rearing (cream).

**Figure 6.**
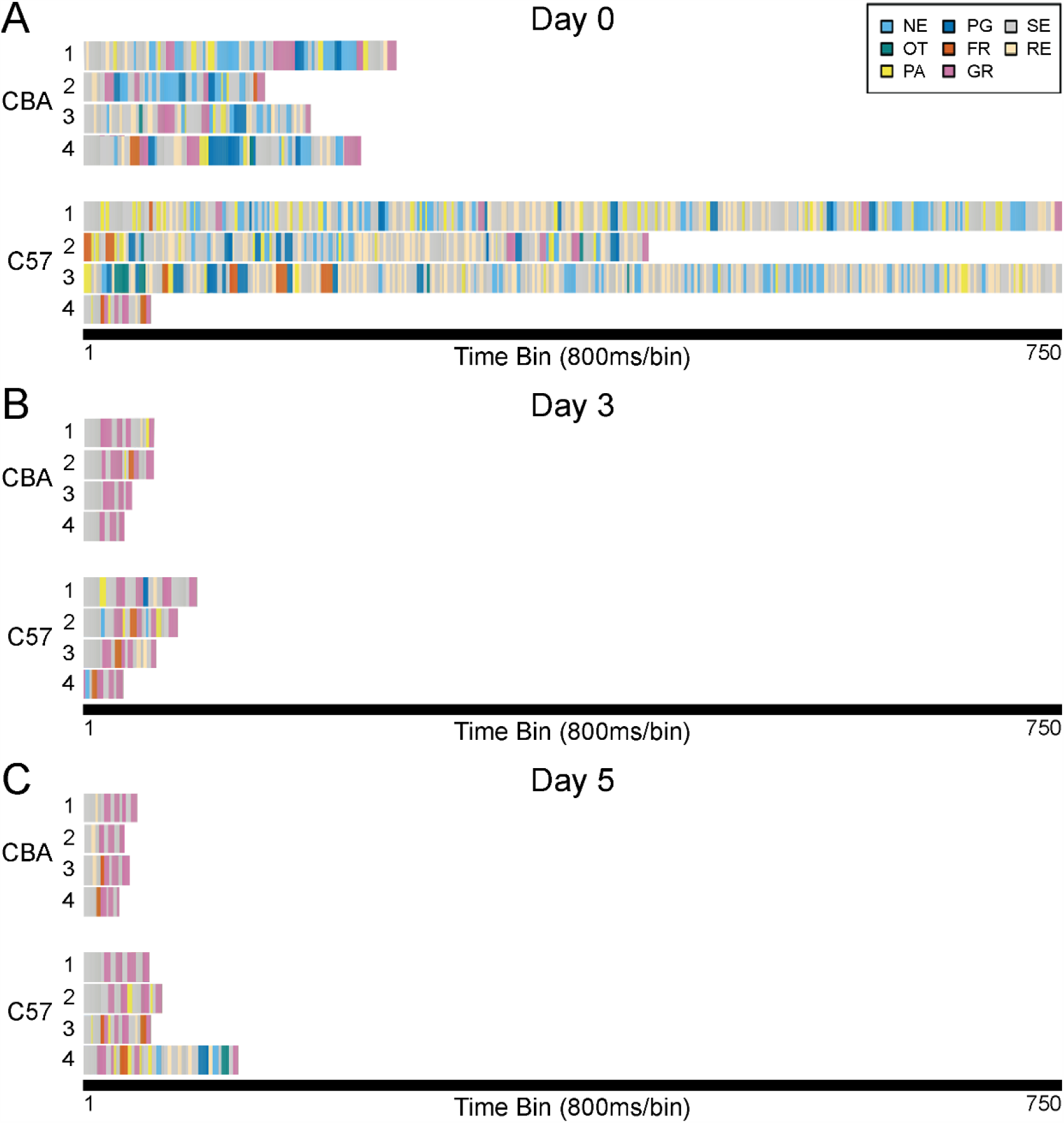
CBA and C57 WT crystallize pup retrieval efficiency by D3 and D5, and exhibit natural variation in sequences of GRMs on D0. **A)** On D0, individual C57 (bottom) performed more GRMs and took longer to retrieve pups than CBA (top). **B-C)** By D3 and D5 (B and C, respectively), mice in both strains had crystallized an efficient pup retrieval sequence. GRMs were coded frame-by-frame during the 10-minute assay. Frames were compressed to 800ms bins for display in the figure. 1^st^ time bin marks the beginning of pup retrieval and ends at the 750^th^ bin (10-minutes). Each row depicts the sequence of behaviors for an individual mouse during retrieval phase. Mouse order is preserved across days to observe patterns and natural variation (example: mouse #1 is the same mouse on D0, D3 and D5). Abbreviations on codes (see Methods for details): NE – nesting (light blue), OT – other (teal), PA – pup approach (yellow), PG – pup grooming (dark blue), FR – failed retrieval (orange), GR – good retrieval (pink), SE – searching (grey), RE – rearing (cream).

**Figure 7.**
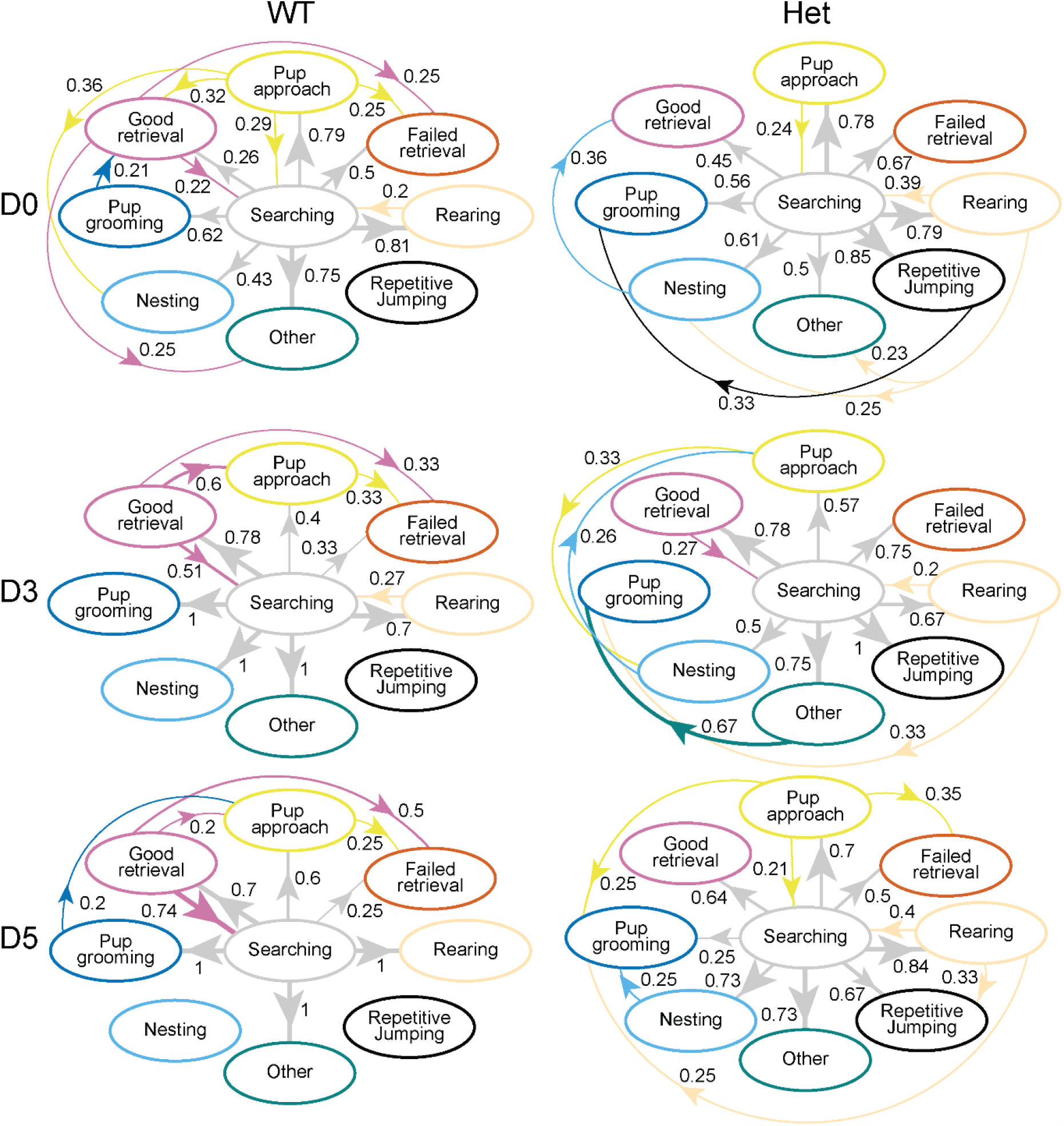
Het exhibit abnormal transition probabilities during retrieval, while WT crystallizes efficient retrieval over days. In WT (left), a predominant pattern of searching (grey) and good retrieval (pink), as indicated by strengthening reciprocal transitional probabilities between the two GRMs, was evident across the three days. In Het (right), this pattern was only observed on D3. The pattern of searching (grey) and pup approach (yellow) was also prominent in WT on D0 and in Het on D0 and D5. Additionally, Het displayed other patterns of transition probabilities between other GRMs as illustrated by the circular connections, while WT displayed limited connections. Transitional probabilities are calculated as lag 1 discrete-time Markov chains. Only probabilities greater than 0.20 are included in this figure. Thicker lines and arrows represent higher probabilities, while thinner lines and arrows represent lower probabilities.

**Figure 8.**
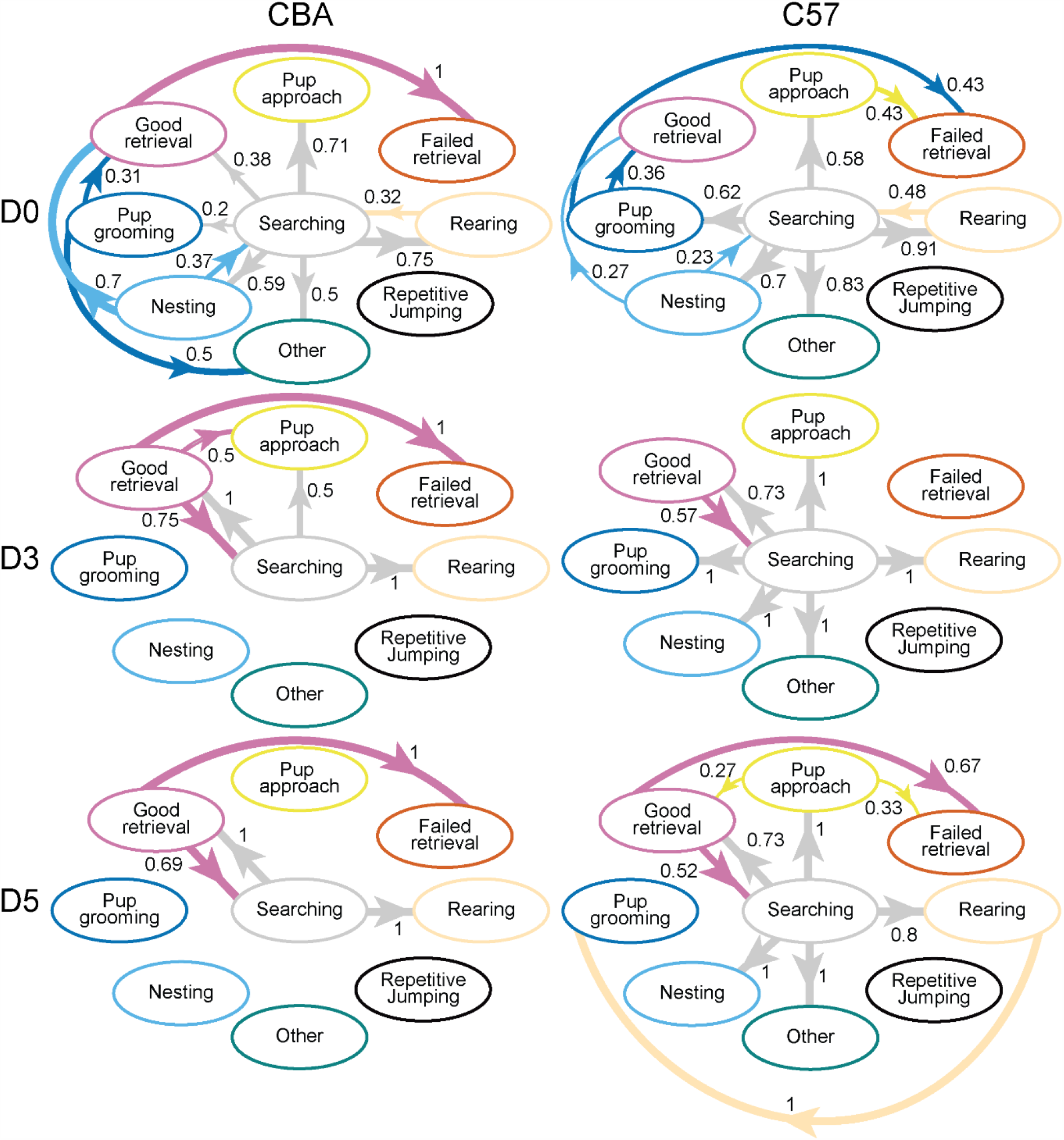
CBA and C57 WT exhibit similar transitional probabilities across GRMs. Although neither CBA (left) nor C57 (right) demonstrated the expected search-to-good retrieval pattern on D0, this pattern became evident in both genotypes at D3 and D5. Transitional probabilities are calculated as lag 1 discrete-time Markov chains. Only probabilities greater than 0.20 are included in this figure. Thicker lines and arrows represent higher probabilities, while thinner lines and arrows represent lower probabilities.

## Results

Previously, we reported that Mecp2^*Heterozygous*^ mice (Het) are inefficient at pup retrieval, as measured by normalized latency index and errors, when the assay was performed on D0, D3 and D5 (Krishnan et al., 2017). As we saw an improvement in Het retrieval efficiency on D3 (Figure 1d, e in Krishnan et al., 2017), we hypothesized that increasing the exposure will lead to better performance. Thus, we performed pup retrieval assay on all six days (D0-D5) here. We found that Het had higher latency index and errors on D0 and D5, compared to WT (Figure 2A, 2B, left). On average, Het had significantly worse performance than WT on both measures (Figure 2A, 2B, right). Thus, performing the pup retrieval assay every day allowed for a better resolution of time course for inefficient retrieval in Het.

As we utilized CBA and C57 strains in this assay, we compared behavioral performance between these WT strains. CBA and C57 surrogates retrieved efficiently according to latency index and error measures (Figure 2C, D). There were no significant differences between the strains across days, though both strains improve significantly on later days, compared to D0. Together, we confirmed our previous findings that Het are inefficient at pup retrieval, with better time resolution. Additionally, we showed that CBA and C57 WT surrogates are efficient at pup retrieval, according to latency index and error measurements.

### Het mice show dynamic changes in specific goal-related movements over days

To capture the dynamic nature of the pup retrieval behavior, beyond the two measures of latency index and errors, we performed frame-by-frame analysis using DataVyu software (Datavyu Team, 2014). Using mouse ethology program developed by Garner et al. (https://mousebehavior.org) as a standard, we marked nine different goal-related movements (GRM) (see Methods) that were consistently observed. We then quantified the time mice spent in each GRM as a percentage of the total time in each different phase (Figure 3). During the habituation phase, where pups and one adult were in the home cage (Figure 1B), Het spent significantly more time engaged in repetitive jumping and significantly less time grooming pups, compared to WT (Figure 3A). No other significant differences were observed in the other GRMs. In the isolation phase, when pups were removed from the home cage (Figure 1B), both WT and Het exhibited fewer repertoire of similar GRMs and spent time similarly for each GRM (Figure 3B). During the retrieval phase (Figure 1B), Het spent significantly more time in repetitive jumping, nesting, rearing and less time in good retrieval, compared to the WT (Figure 3C). During the post-retrieval phase, WT spent more time grooming pups, while Het spent more time with repetitive jumping (Figure 3D), similar to the habituation phase. The individual variability across different animals in time spent per GRM is also evident from this analysis.

Day-by-day analysis of Het compared to WT showed that during the habituation phase (Table 1), Het spent significantly more time in repetitive jumping and pup grooming on D0 and D5, and engaged in more searching on D3. During the retrieval phase, Het spent significantly more time on repetitive jumping and rearing and lesser time in good retrieval on D0, no significant changes in time spent on any GRMs on D3, and, more time on nesting, ‘other’, and significantly lesser time on good retrieval on D5 (Table 3). During post-retrieval phase, Het was similar to WT in time spent on GRMs on D0, significantly more time on repetitive jumping, pup approach and significantly less time on pup grooming on D3, significantly more time in repetitive jumping and lesser time in pup grooming on D5 (Table 4). Together, these results show that no GRM is consistently significantly different in Het, compared to WT, over days and phases, suggesting a context-dependent phenotype in Het, rather than an overall deficit in any particular GRM.

**Table 1.** Habituation phase, WT vs. Het

| GRM | Day | Group 1 |  |  | Group 2 |  |  | z | p |
| --- | --- | --- | --- | --- | --- | --- | --- | --- | --- |
|  |  | WT |  |  | Het |  |  |  |  |
|  |  | N | Median | IQR | N | Median | IQR |  |  |
| Repetitive jumping | 0 | 5 | 0 | 0.010 | 5 | 0.068 | 0.01 | -2.11 | 0.03* |
|  | 3 | 6 | 0.001 | 0.003 | 6 | 0.019 | 0.003 | -1.79 | 0.07 |
|  | 5 | 6 | 0 | 0 | 6 | 0.020 | 0 | -2.59 | 0.01** |
| Nesting | 0 | 5 | 0.050 | 0.172 | 5 | 0.057 | 0.014 | -0.21 | 0.83 |
|  | 3 | 6 | 0.041 | 0.117 | 6 | 0.038 | 0.075 | 0 | 1.00 |
|  | 5 | 6 | 0.024 | 0.064 | 6 | 0.009 | 0.035 | -0.24 | 0.81 |
| Other | 0 | 5 | 0.013 | 0.017 | 5 | 0.013 | 0.010 | -0.31 | 0.75 |
|  | 3 | 6 | 0.038 | 0.084 | 6 | 0.020 | 0.020 | -1.52 | 0.13 |
|  | 5 | 6 | 0.034 | 0.116 | 6 | 0.026 | 0.151 | 0 | 1.00 |
| Good Retrieval | 0 | 5 | 0 | 0 | 5 | 0 | 0 | -0.80 | 0.42 |
|  | 3 | 6 | 0 | 0.013 | 6 | 0 | 0 | -1.35 | 0.18 |
|  | 5 | 6 | 0 | 0 | 6 | 0 | 0 | NaN | NaN |
| Pup approach | 0 | 5 | 0.031 | 0.005 | 5 | 0.032 | 0.024 | 0 | 1.00 |
|  | 3 | 6 | 0.023 | 0.013 | 6 | 0.023 | 0.016 | -0.08 | 0.94 |
|  | 5 | 6 | 0.015 | 0.006 | 6 | 0.018 | 0.017 | -0.08 | 0.94 |
| Pup grooming | 0 | 5 | 0.029 | 0.027 | 5 | 0 | 0 | -2.01 | 0.04* |
|  | 3 | 6 | 0.007 | 0.018 | 6 | 0 | 0 | -1.79 | 0.07 |
|  | 5 | 6 | 0.005 | 0.008 | 6 | 0 | 0 | -2.59 | 0.01** |
| Rearing | 0 | 5 | 0.199 | 0.154 | 5 | 0.230 | 0.078 | 0 | 1.00 |
|  | 3 | 6 | 0.267 | 0.071 | 6 | 0.242 | 0.042 | -0.72 | 0.47 |
|  | 5 | 6 | 0.218 | 0.095 | 6 | 0.206 | 0.108 | -0.08 | 0.94 |
| Searching | 0 | 5 | 0.573 | 0.060 | 5 | 0.635 | 0.082 | -0.63 | 0.53 |
|  | 3 | 6 | 0.554 | 0.079 | 6 | 0.631 | 0.013 | -2.00 | 0.05* |
|  | 5 | 6 | 0.524 | 0.116 | 6 | 0.593 | 0.196 | -0.40 | 0.69 |

Descriptive statistical measures of median, interquartile range (IQR), z and p values for relevant goal-related movements (GRMs), with time course resolution. N= numbers of mice. \* represents significant p values.

**Table 2.** Isolation phase, WT vs. Het

| GRM | Day | Group 1 |  |  | Group 2 |  |  | z | p |
| --- | --- | --- | --- | --- | --- | --- | --- | --- | --- |
|  |  | WT |  |  | Het |  |  |  |  |
|  |  | N | Media<br>n | IQR | N | Media<br>n | IQR |  |  |
| Repetitive jumping | 0 | 5 | 0 | 0.006 | 5 | 0.017 | 0.064 | -1.40 | 0.16 |
|  | 3 | 6 | 0 | 0 | 6 | 0.013 | 0.029 | -1.79 | 0.07 |
|  | 5 | 6 | 0 | 0 | 6 | 0.006 | 0.017 | -1.79 | 0.07 |
| Nesting | 0 | 5 | 0.066 | 0.078 | 5 | 0.215 | 0.212 | -0.63 | 0.53 |
|  | 3 | 6 | 0.063 | 0.035 | 6 | 0.066 | 0.149 | -0.08 | 0.94 |
|  | 5 | 6 | 0.116 | 0.114 | 6 | 0.076 | 0.158 | -0.24 | 0.81 |
| Other | 0 | 5 | 0.020 | 0.006 | 5 | 0.002 | 0.009 | -1.06 | 0.29 |
|  | 3 | 6 | 0.030 | 0.066 | 6 | 0.018 | 0.025 | -0.08 | 0.94 |
|  | 5 | 6 | 0.049 | 0.097 | 6 | 0.035 | 0.075 | -0.56 | 0.57 |
| Rearing | 0 | 5 | 0.197 | 0.032 | 5 | 0.155 | 0.165 | -0.42 | 0.68 |
|  | 3 | 6 | 0.169 | 0.052 | 6 | 0.186 | 0.056 | -0.56 | 0.58 |
|  | 5 | 6 | 0.161 | 0.026 | 6 | 0.220 | 0.154 | -0.72 | 0.47 |
| Searching | 0 | 5 | 0.643 | 0.055 | 5 | 0.643 | 0.093 | -0.21 | 0.83 |
|  | 3 | 6 | 0.689 | 0.093 | 6 | 0.648 | 0.119 | -1.20 | 0.23 |
|  | 5 | 6 | 0.612 | 0.177 | 6 | 0.579 | 0.153 | -0.40 | 0.69 |

Descriptive statistical measures of median, interquartile range (IQR), z and p values for relevant goal-related movements (GRMs), with time course resolution. N= numbers of mice.

**Table 3.** Retrieval phase, WT vs. Het

| GRM | Day | Group 1 |  |  | Group 2 |  |  | z | p |
| --- | --- | --- | --- | --- | --- | --- | --- | --- | --- |
|  |  | WT |  |  | Het |  |  |  |  |
|  |  | N | Media<br>n | IQR | N | Media<br>n | IQR |  |  |
| Repetitive jumping | 0 | 5 | 0 | 0 | 5 | 0.046 | 0.057 | -2.67 | 0.01** |
|  | 3 | 6 | 0 | 0 | 6 | 0 | 0 | -0.83 | 0.40 |
|  | 5 | 6 | 0 | 0 | 6 | 0 | 0.003 | -1.35 | 0.18 |
| Nesting | 0 | 5 | 0.042 | 0.078 | 5 | 0.168 | 0.212 | -1.25 | 0.21 |
|  | 3 | 6 | 0 | 0.035 | 6 | 0 | 0.149 | -0.74 | 0.46 |
|  | 5 | 6 | 0 | 0.114 | 6 | 0.039 | 0.158 | -2.59 | 0.01** |
| Other | 0 | 5 | 0.006 | 0.006 | 5 | 0.021 | 0.009 | -0.63 | 0.53 |
|  | 3 | 6 | 0.011 | 0.066 | 6 | 0.024 | 0.025 | -0.65 | 0.51 |
|  | 5 | 6 | 0 | 0.097 | 6 | 0.045 | 0.075 | -2.05 | 0.04* |
| Failed Retrieval | 0 | 5 | 0.026 | 0.032 | 5 | 0.030 | 0.165 | -0.42 | 0.67 |
|  | 3 | 6 | 0.009 | 0.052 | 6 | 0.013 | 0.056 | 0 | 1.00 |
|  | 5 | 6 | 0.049 | 0.026 | 6 | 0.063 | 0.154 | -0.73 | 0.47 |
| Good Retrieval | 0 | 5 | 0.258 | 0.055 | 5 | 0.013 | 0.093 | -2.51 | 0.01** |
|  | 3 | 6 | 0.436 | 0.093 | 6 | 0.448 | 0.119 | -0.40 | 0.69 |
|  | 5 | 6 | 0.527 | 0.177 | 6 | 0.079 | 0.153 | -2.32 | 0.02* |
| Pup approach | 0 | 5 | 0.206 | 0.135 | 5 | 0.099 | 0.055 | -1.04 | 0.30 |
|  | 3 | 6 | 0.005 | 0.025 | 6 | 0.069 | 0.085 | -1.79 | 0.07 |
|  | 5 | 6 | 0.015 | 0.031 | 6 | 0.135 | 0.184 | -1.79 | 0.07 |
| Pup grooming | 0 | 5 | 0 | 0.099 | 5 | 0 | 0 | -0.77 | 0.44 |
|  | 3 | 6 | 0 | 0 | 6 | 0 | 0 | 0 | 1.00 |
|  | 5 | 6 | 0 | 0 | 6 | 0 | 0.006 | -0.53 | 0.60 |
| Rearing | 0 | 5 | 0.026 | 0.018 | 5 | 0.129 | 0.047 | -2.09 | 0.04* |
|  | 3 | 6 | 0.010 | 0.065 | 6 | 0.002 | 0.022 | -0.34 | 0.73 |
|  | 5 | 6 | 0.024 | 0.037 | 6 | 0.047 | 0.052 | -0.73 | 0.47 |
| Searching | 0 | 5 | 0.403 | 0.120 | 5 | 0.387 | 0.055 | 0 | 1.00 |
|  | 3 | 6 | 0.416 | 0.124 | 6 | 0.416 | 0.030 | -0.08 | 0.94 |
|  | 5 | 6 | 0.392 | 0.242 | 6 | 0.311 | 0.223 | -0.08 | 0.94 |

Descriptive statistical measures of median, interquartile range (IQR), z and p values for relevant goal-related movements (GRMs), with time course resolution. N= numbers of mice. \* represents significant p values.

**Table 4.** Post-Retrieval, WT vs. Het

| GRM | Day | Group 1 |  |  | Group 2 |  |  | z | p |
| --- | --- | --- | --- | --- | --- | --- | --- | --- | --- |
|  |  | WT |  |  | Het |  |  |  |  |
|  |  | N | Media<br>n | IQR | N | Media<br>n | IQR |  |  |
| Repetitive jumping | 0 | 3 | 0 | 0.001 | 3 | 0.148 | 0.148 | -0.43 | 0.67 |
|  | 3 | 4 | 0 | 0 | 4 | 0.009 | 0.006 | -2.43 | 0.02* |
|  | 5 | 4 | 0 | 0 | 4 | 0.008 | 0.008 | -1.97 | 0.05* |
| Nesting | 0 | 3 | 0.289 | 0.280 | 3 | 0.373 | 0.045 | -0.19 | 0.85 |
|  | 3 | 4 | 0.406 | 0.216 | 4 | 0.389 | 0.202 | -1.00 | 0.32 |
|  | 5 | 4 | 0.220 | 0.104 | 4 | 0.326 | 0.082 | -1.19 | 0.24 |
| Other | 0 | 3 | 0.025 | 0.067 | 3 | 0.034 | 0.028 | -0.19 | 0.85 |
|  | 3 | 4 | 0.074 | 0.078 | 4 | 0.087 | 0.159 | -0.64 | 0.52 |
|  | 5 | 4 | 0.120 | 0.082 | 4 | 0.094 | 0.114 | -0.46 | 0.65 |
| Failed Retrieval | 0 | 3 | 0 | 0 | 3 | 0 | 0 | NaN | NaN |
|  | 3 | 4 | 0 | 0.007 | 4 | 0 | 0.005 | -0.11 | 0.92 |
|  | 5 | 4 | 0 | 0 | 4 | 0 | 0 | 0 | 1.00 |
| Good Retrieval | 0 | 3 | 0.019 | 0.012 | 3 | 0.028 | 0.021 | -0.19 | 0.85 |
|  | 3 | 4 | 0.007 | 0.006 | 4 | 0.006 | 0.007 | -0.65 | 0.51 |
|  | 5 | 4 | 0.007 | 0.007 | 4 | 0.012 | 0.021 | -0.65 | 0.51 |
| Pup approach | 0 | 3 | 0.065 | 0.042 | 3 | 0.025 | 0.025 | -0.97 | 0.33 |
|  | 3 | 4 | 0.042 | 0.012 | 4 | 0.080 | 0.064 | -2.10 | 0.04* |
|  | 5 | 4 | 0.044 | 0.022 | 4 | 0.041 | 0.023 | -0.09 | 0.93 |
| Pup grooming | 0 | 3 | 0.010 | 0.043 | 3 | 0.081 | 0.081 | 0 | 1.00 |
|  | 3 | 4 | 0.073 | 0.061 | 4 | 0 | 0 | -2.20 | 0.03* |
|  | 5 | 4 | 0.074 | 0.076 | 4 | 0.007 | 0.023 | -2.29 | 0.02* |
| Rearing | 0 | 3 | 0.101 | 0.011 | 3 | 0.071 | 0.013 | -0.97 | 0.33 |
|  | 3 | 4 | 0.063 | 0.045 | 4 | 0.115 | 0.064 | -1.19 | 0.24 |
|  | 5 | 4 | 0.048 | 0.038 | 4 | 0.050 | 0.042 | -0.82 | 0.41 |
| Searching | 0 | 3 | 0.380 | 0.110 | 3 | 0.240 | 0.015 | -1.74 | 0.08 |
|  | 3 | 4 | 0.281 | 0.081 | 4 | 0.339 | 0.151 | -0.82 | 0.41 |
|  | 5 | 4 | 0.402 | 0.119 | 4 | 0.382 | 0.037 | -0.27 | 0.78 |

Descriptive statistical measures of median, interquartile range (IQR), z and p values for relevant goal-related movements (GRMs), with time course resolution. N= numbers of mice. \* represents significant p values.

Comparing Sur between CBA and C57 strains, we did not observe any significant differences between GRMs in any of the phases, when collapsing across days (Figure 4). The only differences appear in D3 where CBA spends significantly more time nesting and C57 spends significantly more time in searching GRM during habituation (Tables 5-8), showing possible strain-specific differences in habituation phase and not in other phases.

**Table 5.**
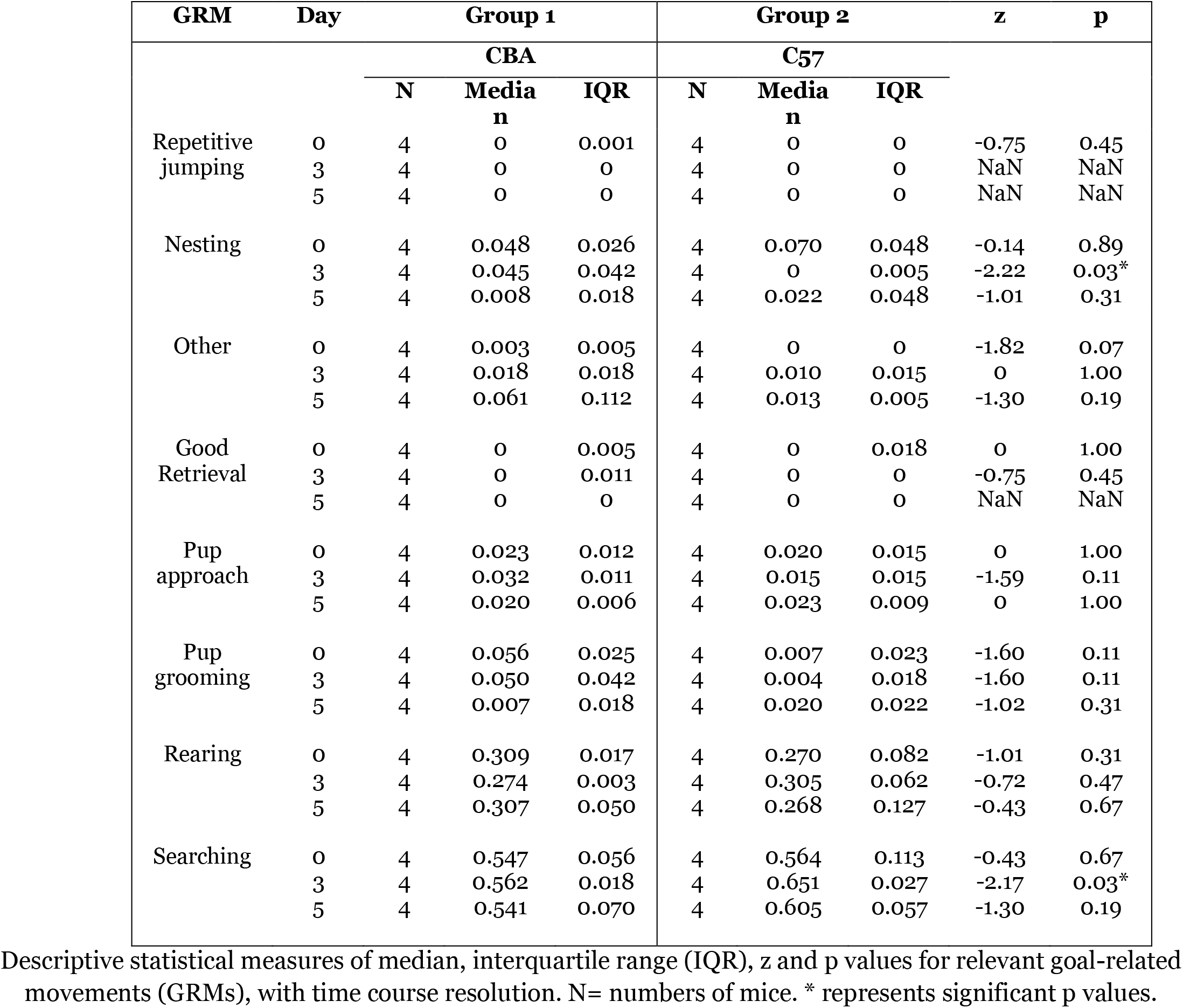
Habituation phase, CBA vs. C57

| GRM | Day | Group 1 |  |  | Group 2 |  |  | z | p |
| --- | --- | --- | --- | --- | --- | --- | --- | --- | --- |
|  |  | CBA |  |  | C57 |  |  |  |  |
|  |  | N | Media<br>n | IQR | N | Media<br>n | IQR |  |  |
| Repetitive jumping | 0 | 4 | 0 | 0.001 | 4 | 0 | 0 | -0.75 | 0.45 |
|  | 3 | 4 | 0 | 0 | 4 | 0 | 0 | NaN | NaN |
|  | 5 | 4 | 0 | 0 | 4 | 0 | 0 | NaN | NaN |
| Nesting | 0 | 4 | 0.048 | 0.026 | 4 | 0.070 | 0.048 | -0.14 | 0.89 |
|  | 3 | 4 | 0.045 | 0.042 | 4 | 0 | 0.005 | -2.22 | 0.03* |
|  | 5 | 4 | 0.008 | 0.018 | 4 | 0.022 | 0.048 | -1.01 | 0.31 |
| Other | 0 | 4 | 0.003 | 0.005 | 4 | 0 | 0 | -1.82 | 0.07 |
|  | 3 | 4 | 0.018 | 0.018 | 4 | 0.010 | 0.015 | 0 | 1.00 |
|  | 5 | 4 | 0.061 | 0.112 | 4 | 0.013 | 0.005 | -1.30 | 0.19 |
| Good Retrieval | 0 | 4 | 0 | 0.005 | 4 | 0 | 0.018 | 0 | 1.00 |
|  | 3 | 4 | 0 | 0.011 | 4 | 0 | 0 | -0.75 | 0.45 |
|  | 5 | 4 | 0 | 0 | 4 | 0 | 0 | NaN | NaN |
| Pup approach | 0 | 4 | 0.023 | 0.012 | 4 | 0.020 | 0.015 | 0 | 1.00 |
|  | 3 | 4 | 0.032 | 0.011 | 4 | 0.015 | 0.015 | -1.59 | 0.11 |
|  | 5 | 4 | 0.020 | 0.006 | 4 | 0.023 | 0.009 | 0 | 1.00 |
| Pup grooming | 0 | 4 | 0.056 | 0.025 | 4 | 0.007 | 0.023 | -1.60 | 0.11 |
|  | 3 | 4 | 0.050 | 0.042 | 4 | 0.004 | 0.018 | -1.60 | 0.11 |
|  | 5 | 4 | 0.007 | 0.018 | 4 | 0.020 | 0.022 | -1.02 | 0.31 |
| Rearing | 0 | 4 | 0.309 | 0.017 | 4 | 0.270 | 0.082 | -1.01 | 0.31 |
|  | 3 | 4 | 0.274 | 0.003 | 4 | 0.305 | 0.062 | -0.72 | 0.47 |
|  | 5 | 4 | 0.307 | 0.050 | 4 | 0.268 | 0.127 | -0.43 | 0.67 |
| Searching | 0 | 4 | 0.547 | 0.056 | 4 | 0.564 | 0.113 | -0.43 | 0.67 |
|  | 3 | 4 | 0.562 | 0.018 | 4 | 0.651 | 0.027 | -2.17 | 0.03* |
|  | 5 | 4 | 0.541 | 0.070 | 4 | 0.605 | 0.057 | -1.30 | 0.19 |

Descriptive statistical measures of median, interquartile range (IQR), z and p values for relevant goal-related movements (GRMs), with time course resolution. N= numbers of mice. \* represents significant p values.

**Table 6.** Isolation phase, CBA vs. C57

| GRM | Day | Group 1 |  |  | Group 2 |  |  | z | p |
| --- | --- | --- | --- | --- | --- | --- | --- | --- | --- |
|  |  | CBA |  |  | C57 |  |  |  |  |
|  |  | N | Media<br>n | IQR | N | Media<br>n | IQR |  |  |
| Repetitive<br>jumping | 0 | 4 | 0.004 | 0.011 | 4 | 0 | 0 | -1.32 | 0.19 |
|  | 3 | 4 | 0 | 0 | 4 | 0 | 0 | NaN | NaN |
|  | 5 | 4 | 0 | 0 | 4 | 0 | 0 | NaN | NaN |
| Nesting | 0 | 4 | 0.188 | 0.115 | 4 | 0.118 | 0.103 | -1.01 | 0.31 |
|  | 3 | 4 | 0.065 | 0.011 | 4 | 0.007 | 0.031 | -1.31 | 0.19 |
|  | 5 | 4 | 0.048 | 0.011 | 4 | 0.080 | 0.129 | -0.14 | 0.89 |
| Other | 0 | 4 | 0.025 | 0.058 | 4 | 0 | 0.001 | -0.83 | 0.41 |
|  | 3 | 4 | 0.046 | 0.054 | 4 | 0.067 | 0.045 | -0.43 | 0.67 |
|  | 5 | 4 | 0.079 | 0.051 | 4 | 0.015 | 0.054 | -1.30 | 0.19 |
| Rearing | 0 | 4 | 0.222 | 0.072 | 4 | 0.191 | 0.035 | -0.72 | 0.47 |
|  | 3 | 4 | 0.201 | 0.071 | 4 | 0.214 | 0.034 | -0.14 | 0.89 |
|  | 5 | 4 | 0.159 | 0.038 | 4 | 0.189 | 0.030 | -0.14 | 0.89 |
| Searching | 0 | 4 | 0.579 | 0.081 | 4 | 0.701 | 0.050 | -1.30 | 0.19 |
|  | 3 | 4 | 0.675 | 0.039 | 4 | 0.677 | 0.034 | 0 | 1.00 |
|  | 5 | 4 | 0.684 | 0.113 | 4 | 0.630 | 0.102 | -0.43 | 0.67 |

Descriptive statistical measures of median, interquartile range (IQR), z and p values for relevant goal-related movements (GRMs), with time course resolution. N= numbers of mice.

**Table 7.** Retrieval phase, CBA vs. C57

| GRM | Day | Group 1 |  |  | Group 2 |  |  | z | p |
| --- | --- | --- | --- | --- | --- | --- | --- | --- | --- |
|  |  | CBA |  |  | C57 |  |  |  |  |
|  |  | N | Media<br>n | IQR | N | Media<br>n | IQR |  |  |
| Repetitive jumping | 0 | 4 | 0 | 0 | 4 | 0 | 0 | NaN | NaN |
|  | 3 | 4 | 0 | 0 | 4 | 0 | 0 | NaN | NaN |
|  | 5 | 4 | 0 | 0 | 4 | 0 | 0 | NaN | NaN |
| Nesting | 0 | 4 | 0.200 | 0.172 | 4 | 0.109 | 0.072 | -1.01 | 0.31 |
|  | 3 | 4 | 0 | 0 | 4 | 0.015 | 0.042 | -1.32 | 0.19 |
|  | 5 | 4 | 0 | 0 | 4 | 0 | 0.019 | -0.75 | 0.45 |
| Other | 0 | 4 | 0.004 | 0.010 | 4 | 0.005 | 0.015 | -0.15 | 0.88 |
|  | 3 | 4 | 0 | 0 | 4 | 0 | 0 | NaN | NaN |
|  | 5 | 4 | 0 | 0 | 4 | 0 | 0.013 | -0.75 | 0.45 |
| Failed Retrieval | 0 | 4 | 0.009 | 0.022 | 4 | 0.020 | 0.034 | -0.74 | 0.46 |
|  | 3 | 4 | 0 | 0.015 | 4 | 0.075 | 0.030 | -1.54 | 0.12 |
|  | 5 | 4 | 0.053 | 0.107 | 4 | 0.025 | 0.079 | 0 | 1.00 |
| Good Retrieval | 0 | 4 | 0.159 | 0.008 | 4 | 0.053 | 0.122 | -1.01 | 0.31 |
|  | 3 | 4 | 0.387 | 0.031 | 4 | 0.311 | 0.072 | -1.01 | 0.31 |
|  | 5 | 4 | 0.397 | 0.034 | 4 | 0.320 | 0.104 | -1.30 | 0.19 |
| Pup approach | 0 | 4 | 0.038 | 0.023 | 4 | 0.046 | 0.033 | -0.14 | 0.89 |
|  | 3 | 4 | 0.004 | 0.010 | 4 | 0.024 | 0.054 | -0.46 | 0.64 |
|  | 5 | 4 | 0 | 0 | 4 | 0.043 | 0.039 | -1.82 | 0.07 |
| Pup grooming | 0 | 4 | 0.083 | 0.054 | 4 | 0.083 | 0.053 | -0.14 | 0.89 |
|  | 3 | 4 | 0 | 0 | 4 | 0 | 0.010 | -0.75 | 0.45 |
|  | 5 | 4 | 0 | 0 | 4 | 0 | 0.018 | -0.75 | 0.45 |
| Rearing | 0 | 4 | 0.056 | 0.042 | 4 | 0.151 | 0.093 | -0.72 | 0.47 |
|  | 3 | 4 | 0 | 0.006 | 4 | 0.011 | 0.036 | -0.50 | 0.62 |
|  | 5 | 4 | 0.052 | 0.046 | 4 | 0.014 | 0.040 | -0.44 | 0.66 |
| Searching | 0 | 4 | 0.390 | 0.054 | 4 | 0.476 | 0.047 | -1.59 | 0.11 |
|  | 3 | 4 | 0.565 | 0.027 | 4 | 0.525 | 0.117 | -1.01 | 0.31 |
|  | 5 | 4 | 0.498 | 0.071 | 4 | 0.531 | 0.099 | -0.72 | 0.47 |

Descriptive statistical measures of median, interquartile range (IQR), z and p values for relevant goal-related movements (GRMs), with time course resolution. N= numbers of mice.

**Table 8.** Post-Retrieval, CBA v. C57

| GRM | Day | Group 1 |  |  | Group 2 |  |  | z | p |
| --- | --- | --- | --- | --- | --- | --- | --- | --- | --- |
|  |  | CBA |  |  | C57 |  |  |  |  |
|  |  | N | Media<br>n | IQR | N | Media<br>n | IQR |  |  |
| Repetitive jumping | 0 | 3 | 0 | 0 | 3 | 0 | 0 | NaN | NaN |
|  | 3 | 4 | 0 | 0 | 4 | 0 | 0 | -0.75 | 0.45 |
|  | 5 | 4 | 0 | 0 | 4 | 0 | 0 | NaN | NaN |
| Nesting | 0 | 3 | 0.266 | 0.159 | 3 | 0.069 | 0.008 | -1.94 | 0.05* |
|  | 3 | 4 | 0.199 | 0.071 | 4 | 0.165 | 0.153 | -0.43 | 0.67 |
|  | 5 | 4 | 0.071 | 0.012 | 4 | 0.113 | 0.113 | -1.01 | 0.31 |
| Other | 0 | 3 | 0.017 | 0.127 | 3 | 0.021 | 0.099 | -0.18 | 0.86 |
|  | 3 | 4 | 0.091 | 0.110 | 4 | 0.169 | 0.085 | -1.30 | 0.19 |
|  | 5 | 4 | 0.491 | 0.097 | 4 | 0.229 | 0.238 | -0.72 | 0.47 |
| Failed Retrieval | 0 | 3 | 0 | 0 | 3 | 0 | 0.002 | -0.87 | 0.39 |
|  | 3 | 4 | 0 | 0 | 4 | 0 | 0 | NaN | NaN |
|  | 5 | 4 | 0 | 0 | 4 | 0 | 0.001 | -0.75 | 0.45 |
| Good Retrieval | 0 | 3 | 0.008 | 0.004 | 3 | 0 | 0.003 | -1.28 | 0.20 |
|  | 3 | 4 | 0.002 | 0.005 | 4 | 0.003 | 0.009 | -0.46 | 0.64 |
|  | 5 | 4 | 0.001 | 0.004 | 4 | 0 | 0 | -0.83 | 0.41 |
| Pup approach | 0 | 3 | 0.046 | 0.021 | 3 | 0.052 | 0.036 | -0.18 | 0.86 |
|  | 3 | 4 | 0.030 | 0.011 | 4 | 0.020 | 0.007 | -1.59 | 0.11 |
|  | 5 | 4 | 0.020 | 0.009 | 4 | 0.013 | 0.003 | -1.01 | 0.31 |
| Pup grooming | 0 | 3 | 0.096 | 0.099 | 3 | 0.006 | 0.015 | -1.59 | 0.11 |
|  | 3 | 4 | 0.064 | 0.071 | 4 | 0.078 | 0.013 | -0.14 | 0.89 |
|  | 5 | 4 | 0.062 | 0.088 | 4 | 0.070 | 0.068 | -0.43 | 0.67 |
| Rearing | 0 | 3 | 0.124 | 0.090 | 3 | 0.208 | 0.049 | -1.24 | 0.22 |
|  | 3 | 4 | 0.101 | 0.036 | 4 | 0.059 | 0.034 | -1.59 | 0.11 |
|  | 5 | 4 | 0.021 | 0.047 | 4 | 0.116 | 0.049 | -1.01 | 0.31 |
| Searching | 0 | 3 | 0.375 | 0.099 | 3 | 0.607 | 0.069 | -1.94 | 0.05* |
|  | 3 | 4 | 0.486 | 0.086 | 4 | 0.409 | 0.089 | -0.43 | 0.67 |
|  | 5 | 4 | 0.324 | 0.120 | 4 | 0.366 | 0.115 | -0.14 | 0.89 |

Descriptive statistical measures of median, interquartile range (IQR), z and p values for relevant goal-related movements (GRMs), with time course resolution. N= numbers of mice. \* represents significant p values.

### Sequences of goal-related movements highlight individual variability across days

In order to fully capture the individual differences in behavior, we plotted the sequences of GRMs during the 10-minute retrieval and post-retrieval phase over days (Figure 5). Individual WT mice improve their retrieval time from D0 to D5. Individual mice displayed varying patterns of sequence on D0, with all WT retrieving all pups within comparable times (Figure 5A). WT #1, #5 and #6 performed searching (SE, grey), followed by good retrieval (GR, pink), followed by pup approach (PA, yellow) and more searching and good retrieval. WT #3 and #4 added pup grooming (PG, dark blue) to that repertoire. Importantly, WT #3, #4 and #5 showed failed retrievals (FR, orange). On D3 (Figure 5B), WT #1 had the longest time to retrieval, while the other WT mice had similar short retrieval times and sequences of GRMs. WT #1, #2, #5 engaged in other (OT, teal) activities in the early part of retrieval while WT #5 was the only one to engage in pup grooming (PG, dark blue). WT #1, #3 and #5 still displayed failed retrieval (FR, orange). On D5 (Figure 5C), WT #1 and #4 had similar patterns with search and retrieved sequences, while WT #3, #5 and #6 displayed failed retrievals. Together, these data show that though on average, WT are efficient at retrieval and exhibit similar sequences, individual mice exhibit natural variation in sequences to pup retrieval on different days. Additionally, WT exhibit a particular sequence motif of “search” directly to “good retrieval” when most efficient in pup retrieval.

On the other hand, individual Het displayed dynamic patterns and variability between mice and on different days (Figure 5). Het #1 improved in retrieval time between D3 and D5, displayed many bouts of pup approach (PA, yellow), nesting (NE, light blue), pup grooming (PG, dark blue), other activities (OT, teal) and repetitive jumping (RJ, black) on D3, which were all minimized on D5. On D0, Het #2 exhibited some repetitive jumping at the beginning of retrieval phase and spent considerable time in the nest in the later parts of the phase. Repetitive jumping was not expressed on D3 or D5. There was improvement in retrieval time with fewer bouts of GRMs not important for pup retrieval. Het #3-6 showed marked improvement in pup retrieval time and sequence from D0 to D3. Interestingly, by D5, Het #3 and #5 increased time in completing pup retrieval and engaged in bouts of sequences involving pup approach, grooming, nesting and other activities. Though the inefficient pup retrieval by Het is known, these results show the extent of variability and the GRM switching in both individual mice and days. The intriguing finding that on D3, many Het exhibit similar patterns of pup retrieval sequence to WT suggest that such a sequence motif towards efficient retrieval is possible for Het, though not exhibited consistently on D0 or D5.

While comparing Sur between CBA and C57, individual C57 on D0 showed more natural variation in retrieval time and in the sequences of GRMs, compared to CBA (Figure 6A). However, all mice in both strains improved in retrieval behavior and sequence on D3 and D5 (Figure 6B-C). The variety of GRMs observed on D0 diminishes by D3 and D5, with marked reduction in nesting (NE, light blue) from D0. Together, these results suggest that both strains exhibit marked natural variation in behavior on D0, when first exposed to pups and pup retrieval; by D3 and D5, the mice have adapted to the pups and retrieval behavior with efficient retrieval process of searching and good retrievals.

### Transition probability analysis reveals efficient pup retrieval sequence over days

Het, in general, display different repertoires during retrieval, evidenced by the different behaviors in sequence, compared to the blocks in colors representing smaller subsets of GRMs in the WT (Figure 5). Thus, to determine the probability of observing a particular GRM after “searching”, we calculated the transition probability (Figure 7). “Searching” features prominently on all days and in both genotypes and thus, forms the central aspect of the graphical representation. In WT (Figure 7, left), on D0, the probability of transition from “searching” to “rearing”, “pup approach”, “pup grooming”, and “other” was very high, with minimal probabilities of transition back to “searching”. On D3, the probability of transition from “searching” to the above GRMs remained high, except for “pup approach”. Other behaviors such as “good retrieval”, “pup grooming” and “nesting” had high probability of showing up after “searching”. Interestingly, transition probability from “good retrieval” to “searching” and “pup approach” also increased in D3. On D5, this behavioral sequence was further solidified as evidenced by high transition probabilities from “searching” to “pup approach”, “good retrieval”, “pup grooming” and “rearing”. Loops from “good retrieval” to “searching” and “failed retrieval” also increased, while “nesting” lost prominent connection from D3. These newer motifs suggest a formation of behavioral sequence that results in efficient retrieval, focusing on the spokes on the top side of the wheel.

On the other hand, Het behavioral motifs do not crystallize over time, and continue engaging the bottom spokes of the wheel (Figure 7, right). On D0, the probability of transition from “searching” to all eight GRMs was high, with minimal reciprocal connections. On D3, “searching” to seven GRMs remained high, with the exception of “pup grooming” which had a higher probability of occurring after “other” behaviors. On D5, the motifs for retrieval efficiency had not emerged, in contrast to WT. These transition probabilities show how differently WT and Het perform their sequences of GRMs during the pup retrieval task.

CBA and C57 showed similar high transition probabilities (Figure 8) from “searching” to “nesting”, “rearing”, “pup approach” and “other” on D0. On D3 and D5, their pattern was simplified with high reciprocal transition probabilities from “searching” to “good retrieval”.

### Non-genetic factors play important roles in efficient pup retrieval

In our experimental design, WT are from a C57BL/6J background but raised by *Mecp2*^*Heterozygous*^ mothers as pups and are typically housed with Het littermates. During the surrogacy experience, they are again co-housed with a Het and a CBA mother. C57 surrogates (in-strain controls), on the other hand, are from C57BL/6J background, raised by C57BL/6J wild type mothers and co-housed with other wild type littermates. Thus, even though WT and C57 surrogates are of the same genetic background, their developmental and social experiences are different. We asked if these non-genetic factors would play a role in how WT performed the pup retrieval behavior (Figure 9).

**Figure 9.**
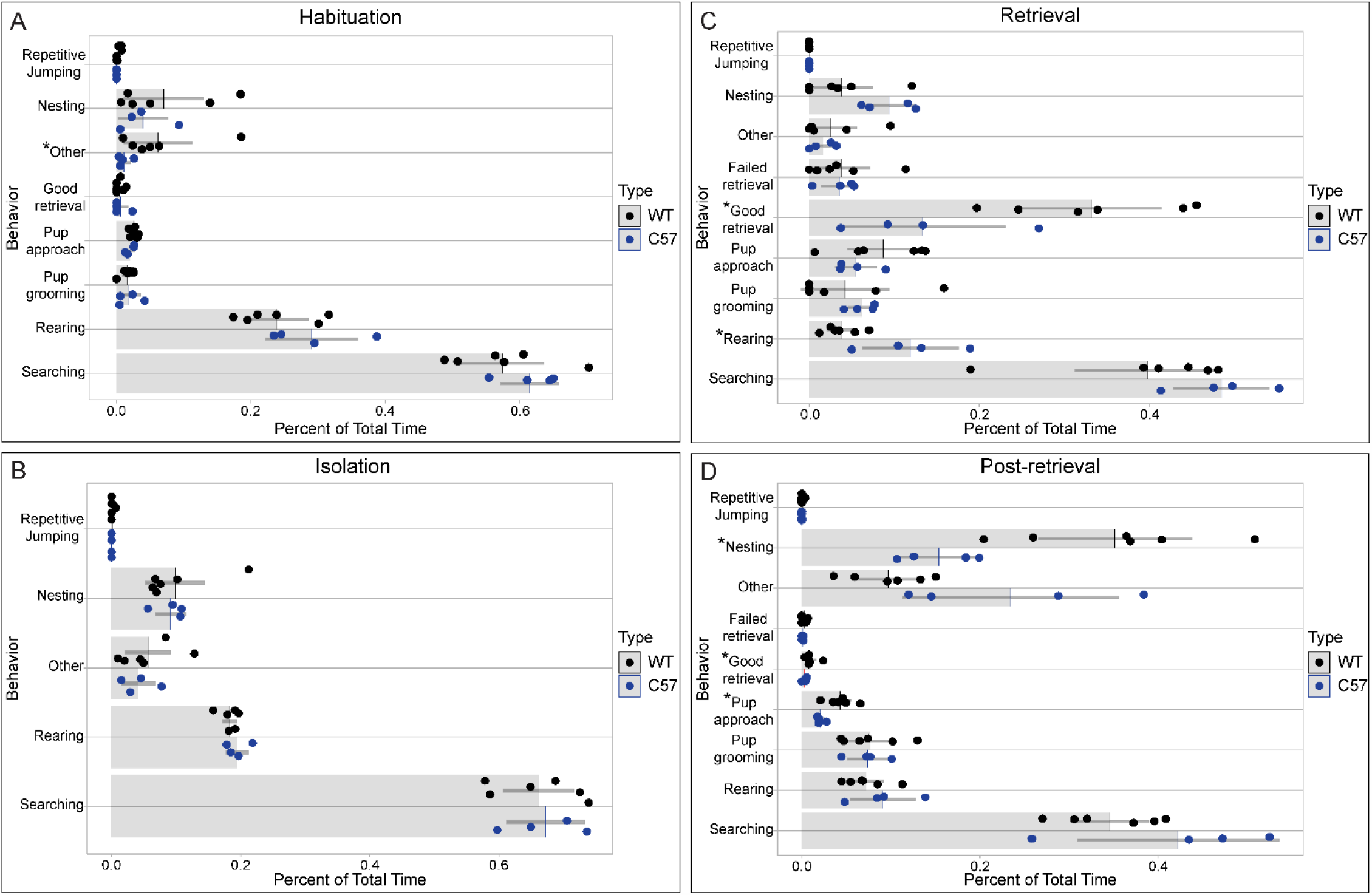
C57 WT and WT (raised by Het mothers) exhibit different GRMs in the presence of pups. **A-D)** While there was no significant difference in any GRMs between C57 (blue) and WT (black) in isolation (B), WT spent significantly more time in ‘other’ during habituation (A), ‘good retrieval’ during retrieval (C), and ‘nesting’, ‘good retrieval’ and ‘pup approach’ in post-retrieval (D). WT also spent significantly lesser time in ‘rearing’ during retrieval (C). These observed differences suggest social experiences affect efficiency in pup retrieval. Light grey bars indicate the mean proportion of time the mice spent in each GRM as a proportion of total time in all GRMs combined. Dark grey lines (whiskers) indicate 95% confidence intervals. Each dot represents the mean across 3 days for each mouse. *Unpaired two-samples Wilcoxon test: *p* < *0*.*05, **p* < *0*.*01*.

Comparing C57 to WT surrogates, significant differences were found in all phases except isolation. More GRMs were affected during post-retrieval and retrieval; particularly, time spent rearing (increased in C57) and good retrieval (increased in WT) during retrieval phase. During post-retrieval phase, WT spent more time in nesting, good retrieval and pup approach, compared to C57. Together, these results suggest that non-genetic factors such as raising conditions and likely social experiences impact efficiency in pup retrieval, in agreement with previous studies (Carlier et al., 1983; Hennessy et al., 1980; Ashbrook et al., 2015; Curley et al., 2010; Shoji and Kato, 2009; Priebe et al., 2005).

## Discussion

Maternal behavior of rodents has been described for over 90 years now (Wiesner and Sheard, 1933). Many different groups have contributed to our knowledge on the hormonal, environmental, epigenetic and neural basis for maternal behavior over the decades. Though much of the early work focused on careful and detailed descriptions of sequences and goal-related movements during maternal behavior in texts, the shift to quantification and graphical representations have ultimately led to a narrower and static view point of this rich behavior with end-point analysis.

Other important considerations are the notable species differences between rats and mice, and quantification of maternal behavior between dams and alloparental females (called “sensitized” females, surrogates, experienced virgins etc.). Much of the early work in this field was primarily done in rat dams. With the explosion of genetic tools since the 1990s, laboratory mouse has emerged as a model system for discovering the genetic and neural basis. The species differences in maternal behavior between rats and mice have not been systematically studied. Unlike naïve virgin female rats, naïve virgin female laboratory mice were shown to “sensitize” and retrieve pups after brief exposures (Leblond, 1940; Noirot, 1972; Gandelman, 1973; Gandelman and Vom Saal, 1975; Carlier et al., 1982). Thus, naïve female laboratory mice were considered to be spontaneously maternal. However, a careful analysis of figures and reading of the literature shows high individual variability in maternal and pup retrieval behavior of naïve female mice, after limited exposures to individual pups for a few days. Particularly, detailed characterization of different aspects of maternal behavior have shown that at least three, one-hour exposures to pups are needed to induce full and/or partial maternal behavior in adult mice (Alsina-Llanes et al., 2015; Stolzenberg and Rissman, 2011). For pup retrieval specifically, cohousing with pups and dam contributes significantly to efficient retrieval in nulliparous mice (Cohen et al., 2011; Marlin et al., 2015; Krishnan et al., 2017; Carcea et al., 2020). Thus, mice still require a sensitization period for efficient expression of maternal behavior, but on a shorter time scale (Alsina-Llanes et al., 2015; Calamandrei and Keverne, 1994; Lonstein and De Vries, 2000; Krishnan et al., 2017). The experimental design and methods of exposure are particularly important in interpreting the underlying neural circuitry, and in determining the role of different factors (sensory information, internal state of the adult, the surrounding environment, specific pup stimuli, hormonal status etc.).

Our study shows in WT surrogates, goal-related movements during pup retrieval behavior over time become stereotyped and crystallized, though individual variations still persist (Figures 5, 7). These results suggest that possibility of differing neural circuitry, with pup stimuli identification, perception and cognitive flexibility in early exposure to familiarity, cognitive inflexibility in the later days, prioritizing speed and efficiency in retrieval. Thus, it is likely that different neural mechanisms across brain regions are involved in this complex behavior. Furthermore, individual variability should be taken into account, especially as the field is rapidly moving towards dissecting specialized neural circuitry for behaviors using optogenetics, chemogenetics and high throughput *in vivo* electrophysiological measurements using dense microelectrode arrays. Additionally, such considerations would lead the field in determining the underlying molecular and cellular mechanisms which give rise to individual phenotypes. After all, it is one brain per animal that is ultimately responsible for the behavior, and not the mean/median across cohorts.

These steps become more important, when viewed from the disorder point of view, in the context of Het. Het behave similarly to WT in the isolation phase, and with minimal GRM differences in habituation and post-retrieval phases; however, they behaved differently during the retrieval phase with high number of GRMs, suggesting context-specific behaviors in the presence of pups. Particularly, Het are affected by tactile interactions with the pups with decreased grooming (involving body/face) and increased pup approach (involving whiskers and face). Both these GRMs suggest atypical tactile processing which could contribute to inefficient pup retrieval. These results are in line with our recent work showing that Het mice have increased and atypical expression of extracellular matrix structures called perineuronal nets, which restrict synaptic plasticity on parvalbumin+ GABAergic interneurons, in specific sub regions of the primary somatosensory cortex (Lau et al., 2020b). Particularly, primary somatosensory cortex regions responsible for tactile sensory perception of whiskers (barrel field cortex), upper lip and jaw are significantly affected in Het. Typically, pup vocalizations trigger suppression of parvalbumin+ GABAergic neuronal responses and a concomitant disinhibition in deep-layer pyramidal neurons in the auditory cortex of WT surrogates (Lau et al., 2020a). In Het surrogates, the increased perineuronal net expression interfere with the synaptic plasticity of parvalbumin+ GABAergic neurons, which ultimately result in lack of disinhibition of the pyramidal neurons (Lau et al., 2020a). It is currently unknown if similar mechanisms also occur in the primary somatosensory cortex, which would ultimately lead to atypical tactile processing in Het. Emerging studies on patients with Rett Syndrome show atypical visual and auditory sensory processing (LeBlanc et al., 2015; Peters et al., 2017; Key et al., 2019), though it is unclear if touch and tactile sensations are affected as well.

Pioneering work in the 1980s and 1990s in rat dams showed that proximal interactions between dams and pups are important for consolidating maternal behavior (Kenyon et al., 1981; Kenyon et al., 1983; Morgan et al., 1992; Stern, 1996) with corresponding changes in the receptive fields of the primary somatosensory cortex of rat dams (Xerri et al., 1994). It is unknown if similar changes occur in mice dams. In WT surrogate mice, we reported an increase in area of the primary somatosensory cortex, in a hemisphere-specific fashion (Lau et al., 2020b). The underlying mechanisms for this area increase after surrogacy experience is unclear.

Repetitive jumping has been reported to be a stereotypy in different strains of mice and in models for neurological disorders (Tanimura et al., 2008; Won et al., 2012; Ryan et al., 2010; Garner et al., 2016). However, WT mice also perform such repetitive jumping in their home cages during maintenance checks (anecdotal reports). The fact that we do not see repetitive jumping in WT during the different phases of pup retrieval behavior, but in Het mainly on D0, suggests novelty or stress-induced stereotypy in some mice.

Though there are differences in GRMs or behavioral repertoire between genotypes and strains in surrogates during pup retrieval behavior, these differences ultimately did not lead to infanticide or pup survival issues in these experiments. Presence and care of the mother for the ∼23 hours when the pup retrieval assay is not being conducted could be a reason for the pup survival. Alternatively, these natural variations in behavioral strategies or “habits for retrieval” are not harmful for pup survival, similar to previous descriptions on mice dams across strains (Champagne et al., 2007).

In conclusion, by systematically characterizing pup retrieval behavior over days with both end-point analysis and dynamics of goal-related movements in surrogate female mice, we set the stage for high throughput video analysis using pose estimation and unsupervised classification, with the intent and long-term goal of determining molecular and cellular mechanisms that contribute to individual behavioral phenotypes.

## Contributions

KK designed the original study. PS and KK performed experiments. PS performed DataVyu analysis. DC performed statistical analysis and generated data visualizations. PS, DC, KK discussed results and wrote the paper.

## Acknowledgements

We would like to thank Dr. Billy Y.B. Lau for finalizing figures in Adobe Illustrator, critical feedback and editing of the manuscript. We would also like to thank other Krishnan lab members for performing the behavior experiments. This work was supported by startup funds from the University of Tennessee – Knoxville (KK), 2019 Office of Undergraduate Research Summer Internship and Nancy Ruth Creery Scholarship to PS.

## Notes

### Competing Interest Statement

The authors have declared no competing interest.

